# Identification of tumor-specific MHC ligands through improved biochemical isolation and incorporation of machine learning

**DOI:** 10.1101/2023.06.08.544182

**Authors:** Shima Mecklenbräuker, Piotr Skoczylas, Paweł Biernat, Badeel Zaghla, Bartłomiej Król-Józaga, Maciej Jasiński, Victor Murcia Pienkowski, Anna Sanecka-Duin, Oliver Popp, Rafał Szatanek, Philipp Mertins, Jan Kaczmarczyk, Agnieszka Blum, Martin Klatt

## Abstract

Isolation of MHC ligands and subsequent analysis by mass spectrometry is considered the gold standard for defining targets for TCR-T immunotherapies. However, as many targets of high tumor-specificity are only presented at low abundance on the cell surface of tumor cells, the efficient isolation of these peptides is crucial for their successful detection. Here, we demonstrate how different isolation strategies, which consider hydrophobicity and post-translational modifications, can improve the detection of MHC ligands, including cysteinylated MHC ligands from cancer germline antigens or point-mutated neoepitopes. Furthermore, we developed a novel MHC class I ligand prediction algorithm (ARDisplay-I) that outperforms the current state-of-the-art and facilitates the assignment of peptides to the correct MHC allele. The model has other applications, such as the identification of additional MHC ligands not detected from mass spectrometry or determining whether the MHC ligands can be presented on the cell surface via MHC alleles not included in the study. The implementation of these strategies can augment the development of T cell receptor-based therapies (i.a. TIL^1^-derived T cells, genetically engineered T cells expressing tumor recognizing receptors or TCR-mimic antibodies) by facilitating the identification of novel immunotherapy targets and by enriching the resources available in the field of computational immunology.

**Significance:** This study demonstrates how the isolation of different tumor-specific MHC ligands can be optimized when considering their hydrophobicity and post-translational modification status. Additionally, we developed a novel machine-learning model for the probability prediction of the MHC ligands’ presentation on the cell surface. The algorithm can assign these MHC ligands to their respective MHC alleles which is essential for the design of TCR-T immunotherapies.

## Introduction

Over the last decade, significant improvements have been made in the isolation and identification of major histocompatibility complex (MHC) bound peptides^1–4^, now allowing the parallel discovery of thousands of MHC ligands in a single experiment. However, different isolation strategies are used for sample preparation, which has a tremendous impact on the isolation of MHC ligands with special biochemical characteristics, such as high hydrophobicity or diverse post-translational modifications (PTMs)^5–7^. Importantly, MHC ligands of high hydrophobicity as well as PTM peptides (e.g. phosphorylation on serine, threonine, or tyrosine) are often more immunogenic compared to their less hydrophobic or unmodified counterparts^8, 9^. Recently, there has been an increase in the awareness of the importance and underrepresentation of cysteinylated MHC ligands as a potential source of immunogenic ligands^10, 11^.

Another crucial step for the identification of relevant targets for T cell immunotherapy and the understanding of T cell immune responses is the correct and precise prediction of antigenic peptides from the entire protein sequences and the subsequent assignment of isolated and/or predicted peptides to the MHC alleles they are actually presented on. For this task, different MHC ligand prediction algorithms have been developed^12–15^ but still, this step is far from being resolved, which leaves open the possibility to improve the definition of MHC ligands for T cell-based immunotherapies. For this reason, to complement the biochemical improvements in target identification, we developed a novel algorithm, termed ARDisplay-I for the prediction of the probability of presentation of MHC class I ligands on the cell surface. The model can also be used for the assignment of such isolated MHC ligands to the corresponding MHC alleles they are presented on. ARDisplay-I was trained on mass spectrometry (MS)-derived data from various sources and enables high generalization to new MHC ligands.

To improve the identification of unmodified, modified and mutated tumor-specific MHC ligands we investigated how different isolation strategies based on the biochemical characteristics and the modification status of a peptide can support the isolation and detection of the respective peptide via liquid chromatography-tandem mass spectrometry (LC-MS/MS). We corroborated previous results on how the use of higher concentrations of acetonitrile can improve the isolation yield of MHC ligands, which can lead to a relative decrease in the isolation quality of the MHC allele with the bound peptide and its preferred anchor amino acid residues. We compared our results for improved MHC ligand isolation with higher concentrations of acetonitrile to the average hydrophobicity of peptides isolated from mono-allelic cell lines^16, 17^, which demonstrated a significant correlation but also showed that the data from mono-allelic cell lines cannot be used to predict which MHC alleles might benefit the most from such isolation strategy. Additionally, we observed trends for the effectiveness of isolating PTM MHC ligands (phosphorylated and cysteinylated) with regard to the hydrophobicity of their modifications. To validate our findings, we then analyzed the effectiveness of different isolation modes for six unmodified, modified (cysteinylated) and mutated tumor-specific MHC ligands quantitatively and qualitatively to highlight the importance of optimized isolation techniques, especially when sample supply is limited.

Overall, this study demonstrated how an optimized isolation strategy improves the isolation of MHC-bound peptides when hydrophobicity and modification status of the peptides are considered. Additionally, we developed a novel MHC ligand prediction algorithm ARDisplay-I, which outperforms the current state-of-the-art and will support not only the definition of new targets for T cell immunotherapies but might also help to improve, for example, predictors of neoantigen fitness^20^, off-target immunotoxicity^18, 19^ and response to immunotherapies, which rely on MHC ligand prediction algorithms^21, 22^.

## Results and Discussion

### The effectiveness of MHC ligand isolation

#### The effectiveness of MHC ligand isolation is positively correlated with the concentration of non-polar solvents used for the separation of peptides and MHC complexes

In a previous study, we have demonstrated that the most influential step in our MHC ligand isolation protocol is the separation of peptides and MHC complexes after loading them onto C18 cartridges^6^. This separation is achieved by specifically eluting the MHC ligands but not the complexes from the C18 material with acetonitrile (ACN). Therefore, we wanted to investigate how the use of a broader range of concentrations of ACN influences the effectiveness of the peptide elution. We hypothesized that not only highly hydrophobic peptides could benefit from the use of high concentrations of ACN but also that very hydrophilic peptides could be better isolated when using very low concentrations of ACN. For our experimental conditions, we chose ACN concentrations of 5%, 20%, 35%, and 50%, and sequential use of all 4 concentrations called the “mix”. Three different cell lines, the multiple myeloma lines JJN3 and LP-1 as well as the lymphoblastic leukemia cell line Nalm-6, were selected for their broad MHC allele repertoires and their expression of MHC class II complexes. MHC complexes of all cell lines were isolated using the MHC-A, -B, and -C specific W6/32 antibodies, and MHC complexes were eluted with trifluoracetic acid (TFA). These eluates were then split into five equal parts and loaded onto C18 cartridges for the five experimental steps. For all ACN concentrations, we saw a clear increase in the number of detectable MHC ligands over all three cell lines and biological replicates (**Fig. 1A**). However, the 5% ACN condition was by far the least effective condition as it could result in an up to 500-fold less effective isolation compared to the elution with 50% ACN (**Fig. 1B**). With 20% ACN, a sufficient isolation of peptides was achieved although MHC ligand yields could still be doubled when using 50% ACN for elution (**Fig. 1A**). Interestingly, although expected to result in the highest peptide yields, the subsequent use of all different ACN conditions did not always result in the highest numbers of isolated MHC ligands (**Fig. 1B**). Still, the subgroups of the isolated MHC ligands would be highly variable between the five different isolation strategies with 10 – 35% of total peptides identified only in one distinct ACN setting (**Fig. 1C**).

**Figure 1.**
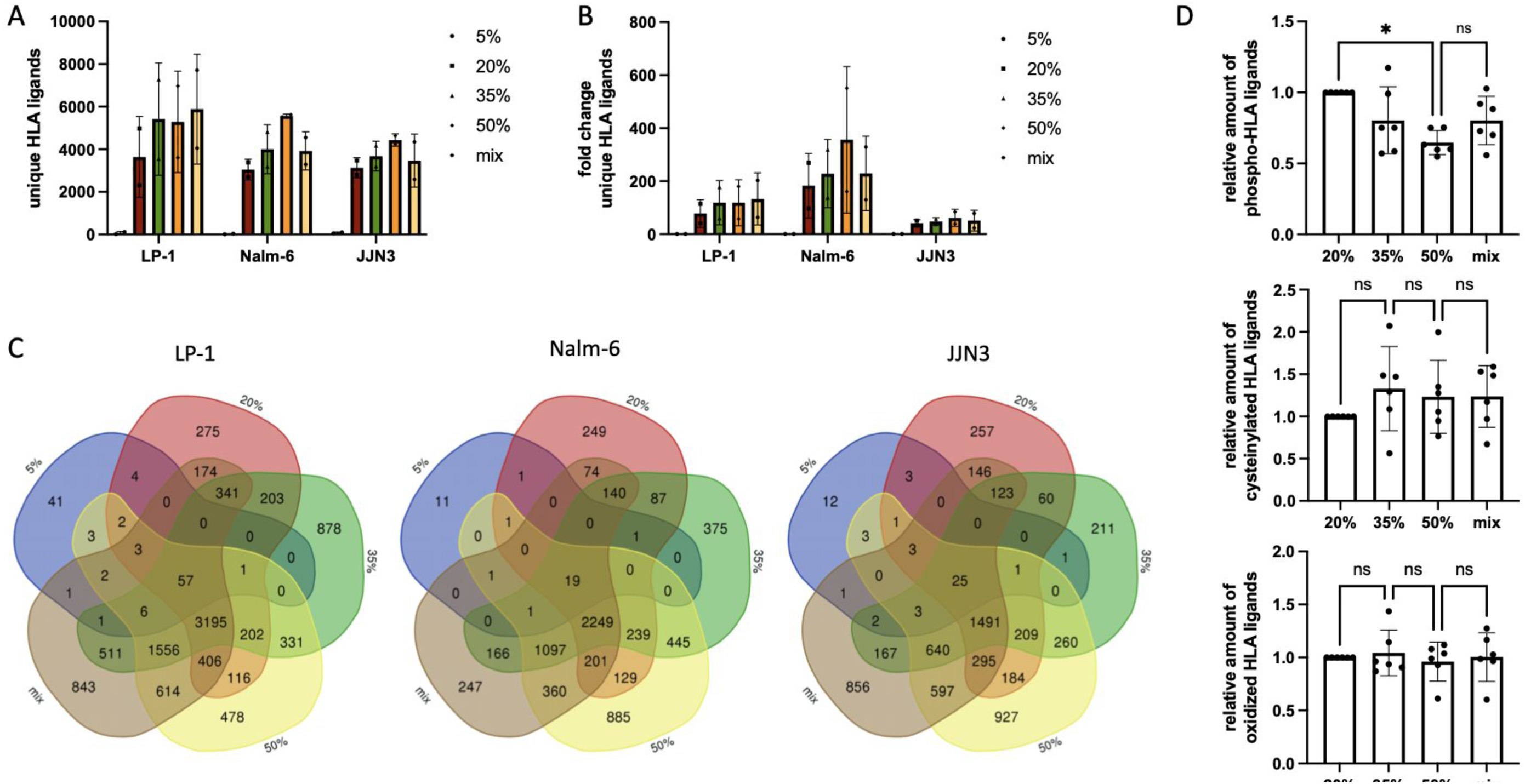
Effect of various conditions of acetonitrile on isolation efficiency of modified and unmodified MHC ligands. **(A)** Unique MHC ligands isolated from the same pool of MHC eluted peptides using various concentrations of acetonitrile **(B)** Relative changes for the yields of unique MHC ligands between different ACN elution conditions in JJN3, LP-1, and Nalm-6 cells. **(C)** Venn diagrams illustrating the overlap of unique MHC ligands in various ACN conditions **(D)** GRAVY scores of different ACN elution conditions in AML14, JMN, and BV173 cells. **(E)** The relative amount of modified MHC ligands isolated from JJN3, LP-1, and Nalm-6 cells. Phosphorylated (top), cysteinylated (middle), and oxidized (bottom). Data were normalized to samples with 5% or 20% ACN. Data are shown for biological duplicates. n.s.=not significant; *, P<0.05.

We then investigated how the different concentrations of ACN affected the recovery of modified MHC ligands. As hypothesized, the biochemical characteristics of the PTMs were essential when the effect on the isolation was analyzed. The addition of a very hydrophilic phosphate-group led to a significantly better isolation of modified MHC ligands in the 20% ACN setting over the 50% ACN setting compared to the isolation of unmodified MHC ligands over all three cell lines (**Fig. 1D top, Suppl. Fig. 1A**). In contrast, cysteinylation of a free thiol group resulted in a trend for better absolute and relative isolation of these MHC ligands in samples eluted with higher concentrations of ACN as the hydrophobicity of these peptides was increased (**Fig. 1D middle, Suppl. Fig. 1B**). Consequently, the oxidation of methionine which does not change the hydrophobicity of a peptide considerably, had no relevant effect on the number of MHC ligands isolated between the different ACN subgroups (**Fig. 1D bottom, Suppl. Fig. 1C**). Similar to the results for unmodified peptides regardless of the type of modification, MHC ligands specific for each ACN concentration were observed although trends for higher numbers in the higher ACN conditions were observed for cysteinylated ligands and in the lower ACN conditions for phosphorylated ligands exemplified for the LP-1 cell lines (**Suppl. Fig. 1D**).

Overall, we demonstrated how higher ACN concentrations correlate with higher numbers of identified MHC ligands although a minimum of 20% seems necessary for sufficient isolation. Additionally, we showed how the amount of ACN used also affects the effectiveness of isolating PTM MHC ligands in dependence on their biochemical characteristics.

### How acetonitrile concentrations facilitate the identification of MHC alleles

#### A wide range of acetonitrile concentrations facilitates the identification of MHC alleles that benefit from the use of high or low acetonitrile concentrations

As the effect of ACN on MHC ligand isolation depends on the MHC allele they are presented on and the resulting anchor amino acids^6^, we next wanted to broaden the knowledge about MHC alleles that benefit from extended or limited use of ACN during the isolation process. For this purpose, we analyzed the 17 different MHC class I alleles expressed on JJN3, LP-1 and Nalm-6 cells, and investigated how the isolation of MHC-specific ligands changed between the highly different ACN concentrations. First, all identified MHC ligands were categorized according to the MHC alleles they are predicted to be presented by, as defined by netMHCpan 4.1 and ARDisplay-I, accordingly, and we then calculated the fractions of MHC ligands relative to the total number of MHC ligands per sample. These results were normalized to the 5% ACN samples, which although low in total MHC ligand yields, provided clear preferences for peptides with low hydrophobicity. Finally, these fractions were compared to the increase of MHC ligands over different ACN conditions (**Fig. 2A-C, Suppl. Fig. 2A-C**). Using this analysis, we were able to define MHC alleles A*02:01, B*08:01, B*14:02, B*15:01, B*18:01, C*04:01, C*07:01, C*07:02, C*08:02 as alleles that benefit from the use of higher ACN concentrations with the strongest increases for A*02:01 and B*08:01 in line with previous results^6^. In contrast, A*01:01, A*03:01, A*33:01 and most profoundly A*30:01 demonstrated relative decreases in MHC ligand isolations (**Fig. 2A-C**) although it must be noted that all MHC alleles, in general, showed higher numbers of isolated MHC ligands with higher concentrations of ACN (**Fig. 2D-F**).

**Figure 2.**
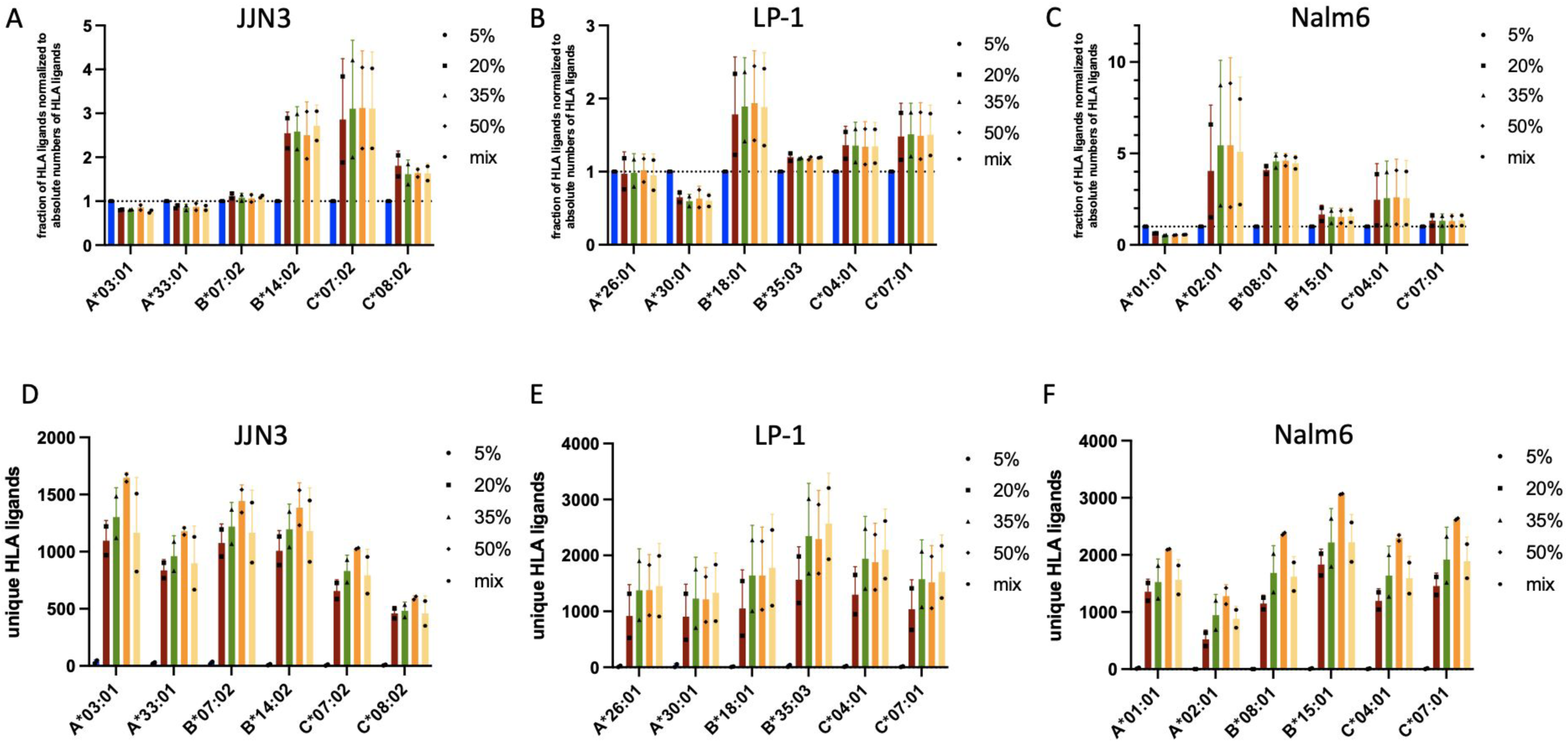
Absolute and relative changes in unique MHC ligands by MHC allele. **(A-C)** Relative changes for the yields of unique MHC ligands between different ACN elution conditions for different alleles expressed in JJN3 (A), LP-1 (B), and Nalm-6 (C) cells. Results are normalized to a 5% ACN setting and relative to the total number of MHC ligands per condition. MHCs ligands were assigned to respective alleles by 1/ netMHCpan using a 2% rank cutoff and 2/ ARDisplay-I with a 0.5 threshold for presentation probability. **(D-F)** Absolute numbers of unique MHC ligands per MHC allele. All experiments were performed in biological duplicates.

To investigate whether MHC ligands of a particular allele would benefit from the use of high concentrations of ACN we then compared the increase of MHC ligands per allele with the average hydrophobicity of peptides eluted from mono-allelic cell lines as these data provide the most reliable resource for binders of a specific MHC allele. Of note, when hydrophobicity scores were calculated for mono-allelic data via the GRAVY score calculator we doubled the values for amino acids at anchor positions as we had previously demonstrated the strong dependence of the MHC ligand isolation from the biochemical characteristics of these amino acids. Additionally, when the increase of MHC ligands per allele was calculated, we used only the results from the 5% ACN and 50% ACN conditions and MHC ligands meeting more stringent prediction criteria to focus mostly on peptides that are uniquely assigned to a single MHC allele rather than multiple alleles, decrease the risk of the inappropriate match and to enable comparability with mono-allelic data. A clear positive and highly significant correlation (R2=0.6080, p=0.0004) was found for the increase of MHC ligands from 5% to 50% conditions with the average hydrophobicity of peptides isolated from mono-allelic cell lines (**Fig. 3**). Still, although highly significant, the correlation was not strong enough to deduct predictions about the usefulness of higher concentrations of ACN for specific MHC alleles.

**Figure 3.**
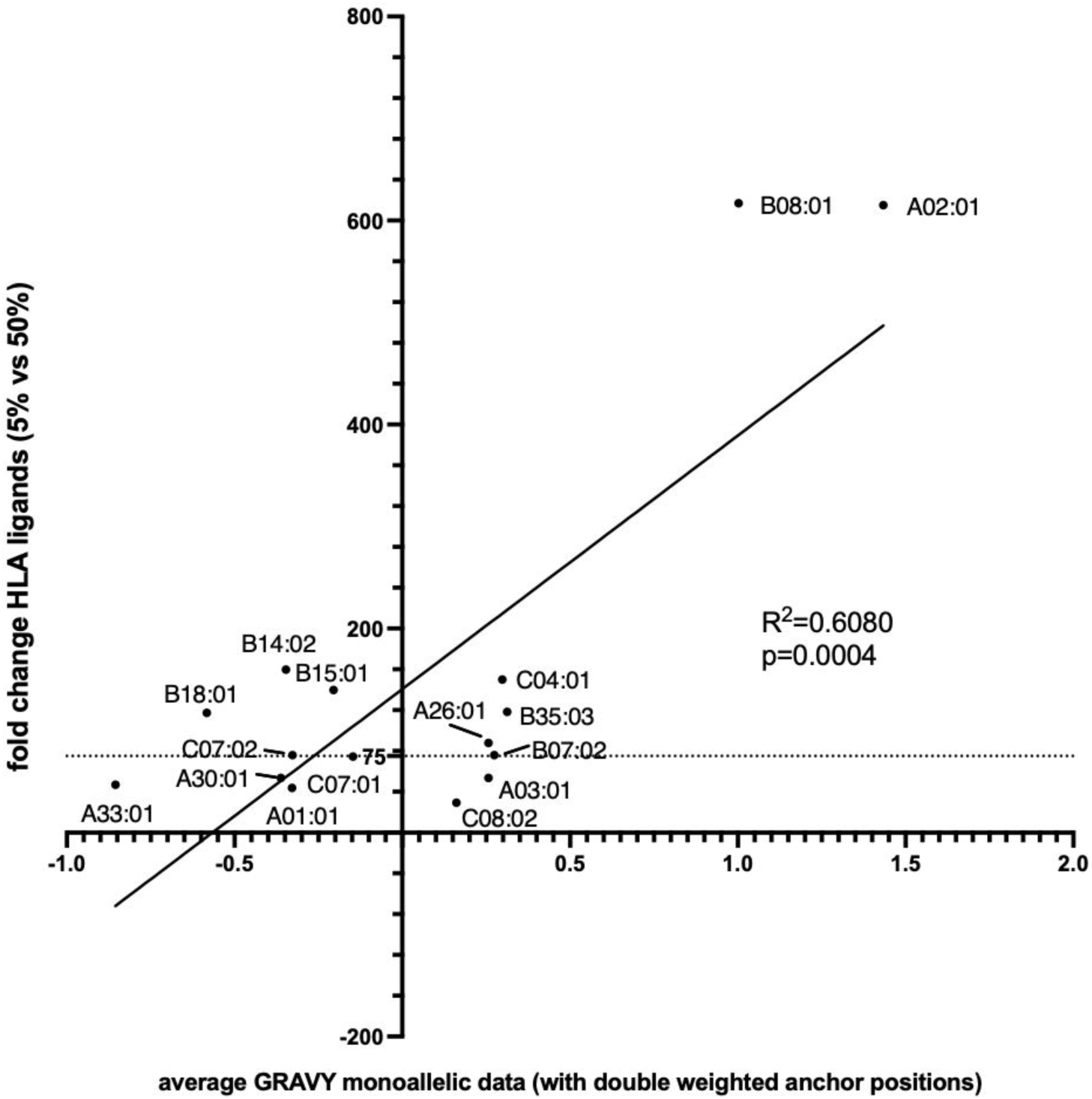
Correlation of the increase of MHC ligand yields per MHC allele and the average hydrophobicity of peptides eluted from mono-allelic cell lines. Average GRAVY scores were calculated for peptides reported by Sarkizova et al. and Abelin et al. from monoallelic cell lines^16, 17^. GRAVY scores of amino acids at anchor positions were doubled. For better comparability, only 9mers were used. Data were correlated with experimental data of 5% and 50% ACN conditions. Peptides isolated in these conditions with a predicted %rank of <0.05 were considered binders.

To enable the analysis on single-allelic data, we selected all observations with netMHCpan %rank scores below 0.05 instead of 2, and for ARDisplay-I, we chose the pHLA pairs with presentation probability higher than 0.5. Even though the threshold used for netMHCpan predictions is much lower than the standard one, the number of MHC ligands obtained is 12.4% higher than by using ARDisplay-I (**Fig. 4G**). When comparing the number of MHC ligands with the distinction of various ACN conditions, the distributions were similar between the two models albeit ARDisplay-I consistently returned a bit smaller quantities than netMHCpan (**Fig. 4A-C**). The differences between the individual MHC alleles were more visible, for example, ARDisplay-I predicted far fewer MHC ligands being presented by HLA-A01:01, A26:01, A33:01, B18:01, B35:03, C04:01, C08:02 while for C07:01, C07:02 it was just the opposite (**Fig. 4D-F**). Since the differences between the results of the two methods are significant, we investigated which prediction is more accurate when validated using a collection of single-allelic data obtained from Sarkizova^17^ and the IEDB database (http://www.iedb.org/) (**Fig. 4G-H**). For the JJN3 and Nalm6 cell lines, more than 75% of overlapping MHC ligands, ie. predicted simultaneously by both algorithms, are experimentally confirmed. The number of MHC ligands predicted exclusively by netMHCpan is much higher than those obtained using ARDisplay-I and rarely any of those is confirmed by single-allelic data. For the three analyzed cell lines we observe 5.9 to 21.2 times higher numbers of experimentally confirmed MHC-matched ligands for ARDisplay-I, as compared to netMHCpan.

**Figure 4.**
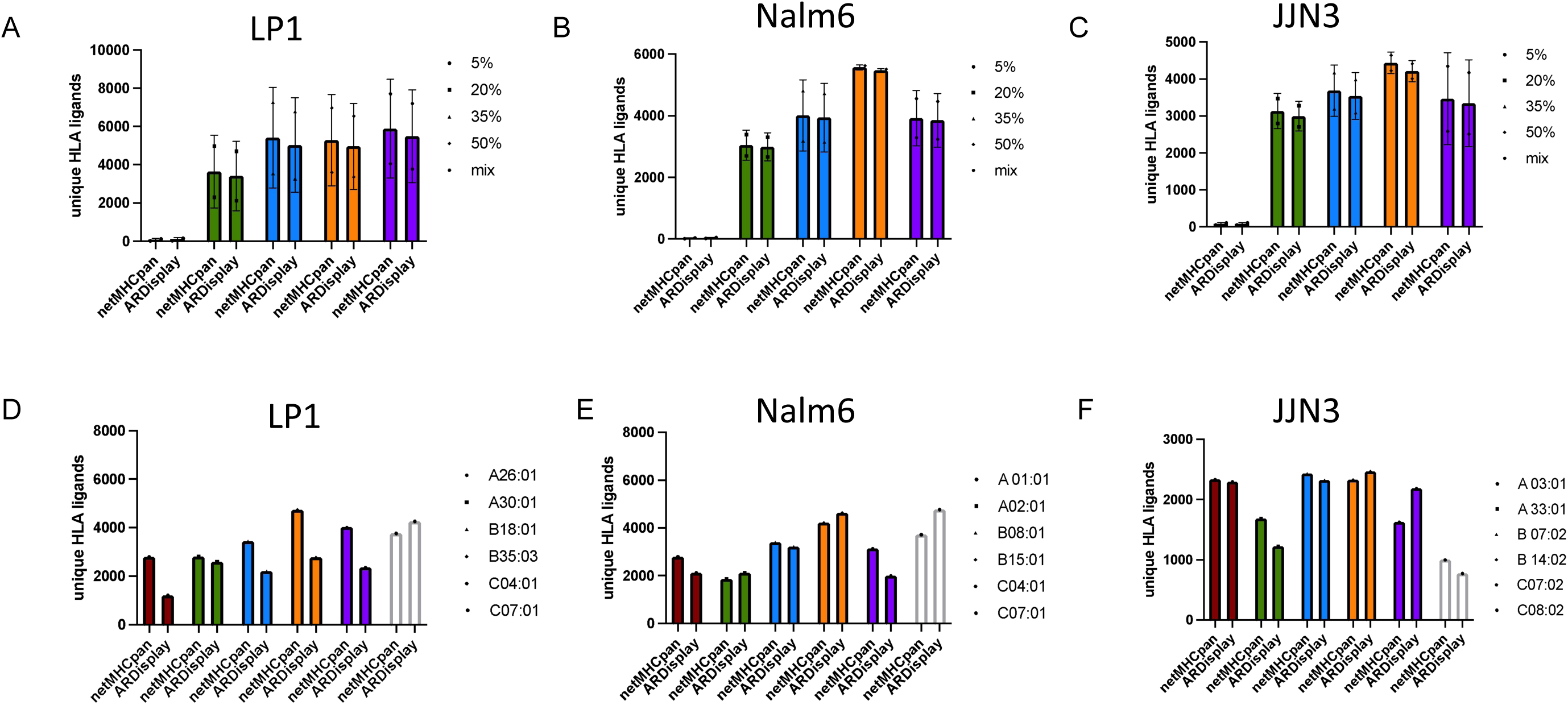
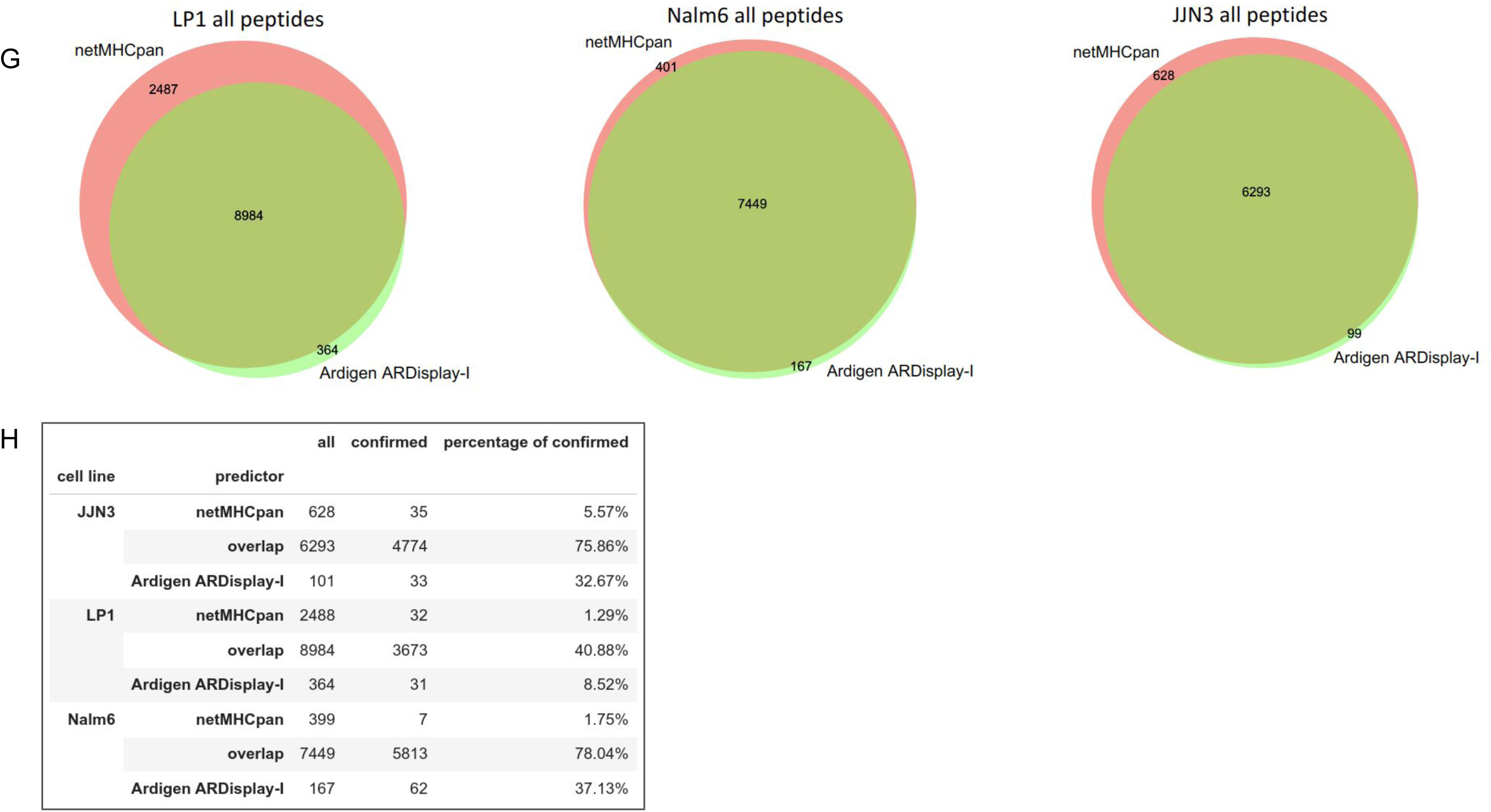
Comparison of netMHCpan with ARDisplay-I in matching MHC ligands with MHC alleles. (A-F) Absolute numbers of unique MHC ligands between different ACN elution conditions and for different MHC alleles expressed in consecutive cells, ie. JJN3, LP-1, and Nalm6 (G) The Venn diagrams with numbers of pMHC pairs predicted by the respective algorithm, or both, as presented on the cell surface (H) Table showing the percentage of predicted observations that were confirmed using single-allelic data sets.

Overall, these data might indicate that for the improved isolation of a specific MHC ligand, the individual biochemical characteristics must be considered rather than its assignment to a specific MHC allele.

### Optimized identification of tumor-specific MHC ligands

#### Identifications of tumor-specific MHC ligands are optimized by considering hydrophobicity and PTMs

As the ideal isolation conditions might not be deduced simply by the MHC allele a peptide is bound to, we now wanted to illustrate that the actual hydrophobicity of a peptide predicted by the GRAVY score as well as the presence or absence of PTMs can support the optimized isolation of certain MHC ligands. First, we re-analyzed our previously published dataset on AML14 cells under the different ACN conditions (30%, 40%, and 50%) with cysteinylation as variable modification as we now have highlighted the importance of this PTM.

Strikingly, two MHC ligands with cysteinylations, derived from cancer germline antigens (CCNA1 and GTSF1), were identified of which the peptide SLSEIVPCysL showed high hydrophobicity and in consequence was only identified in settings with higher concentrations of ACN (40% and 50%) (**Fig. 5A**). We then quantified the signals of several peptide precursors from cancer germline antigens of different hydrophobicity as well as an NRAS Q61L derived neoepitope (**Fig. 5A**) and were able to show profound differences under different ACN isolation conditions (**Fig. 5B**) so that always one experimental setting clearly showed optimal isolation conditions. Although some of the peptides were still found in all investigated samples other peptides were not detected due to non-isolation of the peptide precursors under the 30% ACN condition that would be considered the standard for MHC ligand isolation.

**Figure 5.**
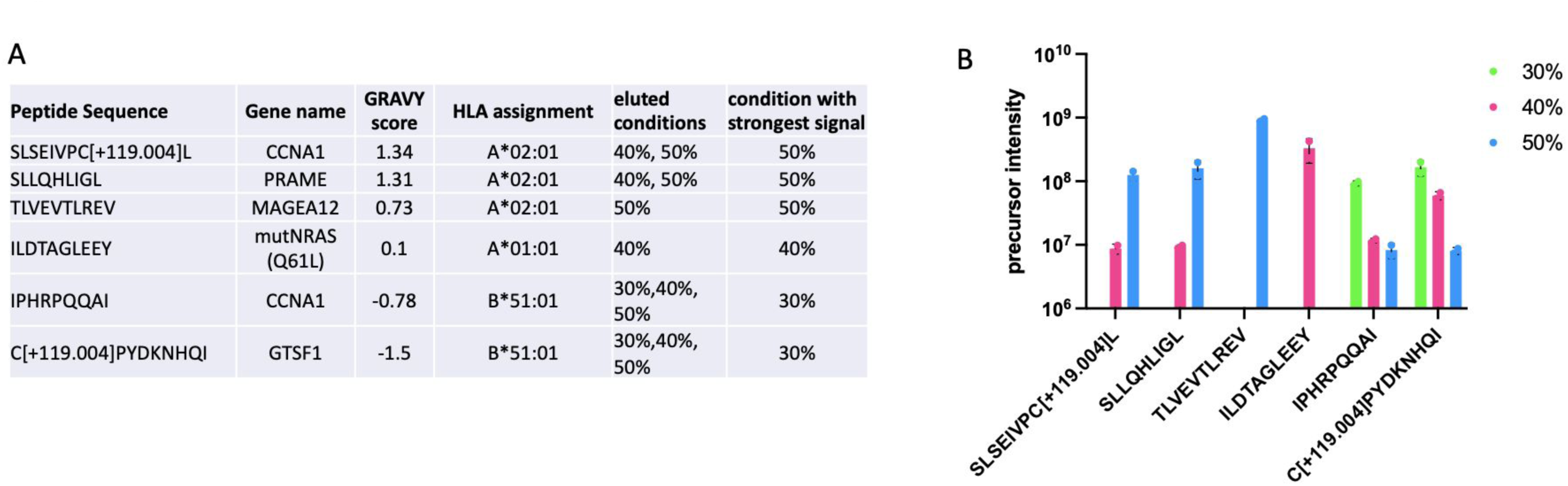
Quantitative changes of tumor-specific MHC ligand under different elution conditions of acetonitrile. **(A)** Table with cancer germline antigen-derived MHC alleles with and without modifications as well as NRAS-derived neoepitope. **(B)** Quantitative changes for individual MHC ligands under different elution conditions of acetonitrile using Skyline^27^.

### ARDisplay-I, HLA^2^ class I ligand presentation model

#### Why use *in silico* predictions?

Even though MS experiments are getting more accessible and easier to perform, they are still unable to overcome some naturally occurring difficulties^3^. While using mono-allelic cell lines, one needs to be aware that typically, retrovirally transduced B-cells are used for such experiments^16, 17, 22^. There are two major drawbacks associated with this approach; 1/ the protein profile of these cells differs substantially from the remaining cells building other tissues, and 2/ while the expression level of the remaining MHC alleles is drastically reduced, it is still possible that residual levels of these alleles are still present^23^, which may lead to peptide presentation on the cell surface, not by the MHC of interest.

These drawbacks can be overcome by using data acquired from healthy donors or cancer patients. However, it should be noted that each patient has its own particular set of HLA-I alleles (up to six different types of canonical HLA-I types, i.e. A, B, and C), and matching peptides with their respective HLA allele is usually based solely on allele-binding predictors^1, 16^. Furthermore, HLA genes are highly polymorphic - as of December 2022 (see https://hla.alleles.org/alleles/index.html), there are almost 25,000 known and described canonical HLA-I alleles (more precisely 7,712 HLA-A, 9,164 HLA-B, and 7,672 HLA-C). Of course, some alleles differ in places unrelated to the peptide binding, referred to as the binding groove, but still, the number of possible combinations is enormous. Namely, accounting for HLA homozygosity (where an individual expresses two identical HLA haplotypes) the total number of possibilities can be estimated based on the formula for sampling with replacement at almost 30,000,000, 42,000,000 and 30,000,000 peptide-HLA (pHLA) pairs for a given peptide and pairs of HLA-I alleles of type A, B and C, respectively. Since those samplings are independent of each other, in total, it is possible to obtain approximately 3,67*10^22^ possible sets of 6 HLA-I alleles.

#### Biological context

Only a tiny fraction of protein fragments are presented on the cell surface and thus can be targeted by T-cells. Accurate prediction of such peptides using *in silico* methods is possible only when taking into account the entire antigen presentation process^17^. As machine learning approaches based on neural networks require extensive training sets, integrating data from a variety of high-quality sources is an essential step to achieving models with high performance. In ARDisplay-I, we achieve this by incorporating MS-eluted ligands together with a customized approach to data processing that includes the generation of negative examples (decoys), ie. pHLA pairs that for various reasons are not displayed on the cell surface.

#### A predictive model for HLA ligands presentation

It was repeatedly shown that models based solely on binding affinity (BA) measurements are insufficient for accurate detection of HLA ligands and models leveraging MS data can lead to higher accuracy, as these models account for the entire antigen processing and presentation pathway^17^. In our approach, we incorporated a biologically aware approach to decoys generation by introducing a set of artificial negative examples from the remaining fragments of proteins matched uniquely to the HLA ligands detected by MS. Additionally, we combined results from BA assays with MS eluted ligands by limiting the training decoys to pHLA pairs with BA below 2000 nM as predicted by MHCflurry^13^.

#### Benchmark and comparison of various HLA ligand prediction models

Currently, there are many computational models that identify the hidden motifs related to pMHC binding and presentation on the cell surface^12–15, 24^. Even though BA predictors have been thoroughly tested (as they were available much earlier than eluted ligand predictors), it seems that they do not generalize well to the task of eluted ligand prediction^17, 25^.

To reliably compare the effectiveness of the different algorithms for eluted ligand (EL) and BA prediction, we used the MS data generated in the present study (described in the previous sections), ie. three different cell lines Nalm6, LP1 and JJN3 expressing 17 distinct HLA class I alleles. We obtained more than 32,000 HLA ligands presented on the cell surface. MS results do not directly provide information about peptides that exist within a cell but are not displayed on its surface via HLA molecules, hence we also generated a set of artificial negative examples from the remaining fragments of original proteins that match uniquely to the HLA eluted ligands.

Consistently with previous results^26^, we showed that using BA predictors is not sufficient to accurately predict HLA ligands and the performance of such algorithms is 2-3 times lower than that of eluted ligand predictors offered by the same groups (**Fig. 6A**). Moreover, ARDisplay-I achieves much higher average precision than the remaining methods - over 2.4 times higher than netMHCpan v4.1 (EL) and over 4.2 times higher than MHCflurry (EL). Additionally, the differences have very high statistical significance (as seen from the non-overlapping confidence bands of the two models). Here, it is also worth noting that the metrics obtained in our benchmark have a substantial gap to the theoretical values for a perfect model (ie. the average precision of 1 for a perfect classifier), which indicates that the benchmark described herein explores a more difficult scenario than the validation schemes typically proposed by other groups^12, 13^. This might be related to our approach to artificial decoys generation and/or a higher ratio of negative to positive observations in our test set.

**Figure 6.**
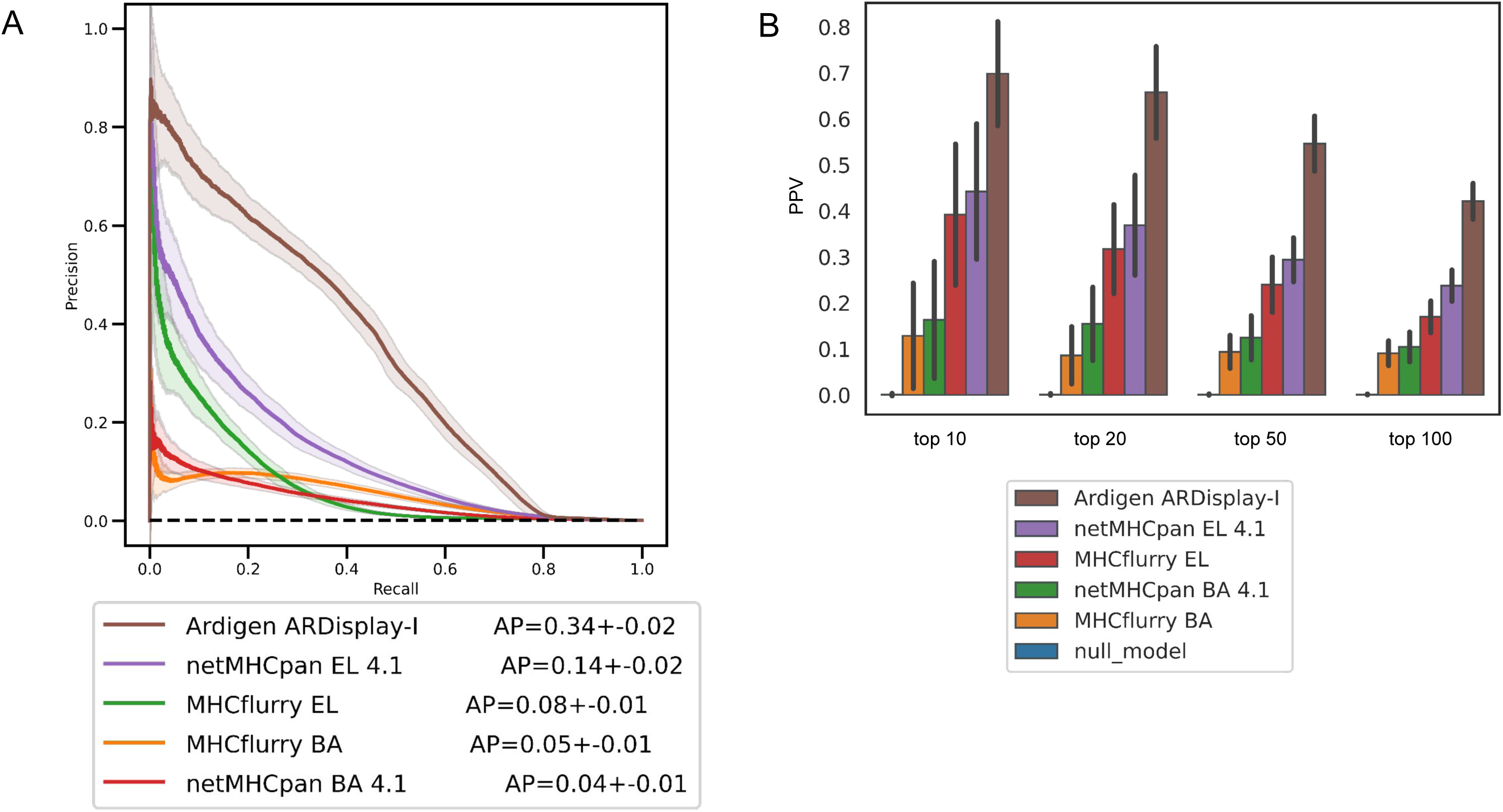
Comparison of Ardigen ARDisplay-I performance with the performance of selected methods. **(A)** Precision-recall curve (PR curve) for the selected methods - see *Materials and Methods* sections for more details. Random resampling of observations from all three cell lines combined was performed 100 times to assess the variability of the metric, ie. estimate the average performance and its confidence band. For each iteration, a set of 500 positives and 500k negatives (x1000 number of positives) was randomly selected. **(B)** Precision (also PPV) for four thresholds, ie. top-10, 20, 50, & 100 MHC ligands selected by each method. The same sampling procedure has been applied as in **(A)**.

To compare the performance of each method, we show the precision (positive predictive value, PPV) for the cases when the top-10, -20, -50 and -100 peptides are selected with each model. Within the top-100 HLA ligands predicted by each algorithm, ARDisplay-I is shown to detect 43+/-0.05 of the presented peptides, whereas netMHCpan v4.1 (EL) detects 23+/-0.04 and MHCflurry (EL) detects 17+/-0.04, and the BA models detect around 10 and 9, respectively. Using BA predictions leads to 2-3.2 times lower precision than using eluted ligand predictions from netMHCpan 4.1 (**Fig. 6B**). Similar conclusions can be reached after analyzing the MHCflurry predictions although in this case, the difference is not as big.

## Conclusions

The isolation and definition of MHC ligands which might serve as targets for T cell-based therapies can be already biased during the isolation process of MHC-bound peptides. As we have demonstrated before, the hydrophobicity of peptides is an important factor that must be taken into account^6^. Here, we broaden the knowledge about alleles that benefit from more rigorous isolation strategies and demonstrate that PTMs also contribute to the detection or non-detection of MHC ligands under specific isolation conditions. Ultimately, optimized isolation conditions for unmodified, modified and mutated MHC ligands were observed and suggest that the knowledge about these optimized settings might facilitate the isolation and definition of T-cell therapy targets.

To complement these biochemical improvements, we developed a novel HLA ligand prediction model, termed ARDisplay-I, and compared its predictive performance with other commonly used methods. In this benchmark, we used the dataset generated in the present study, which enables an unbiased comparison, as the dataset was not used in training any of the benchmarked models. We found that ARDisplay-I achieves over 2.4 times higher average precision than netMHCpan 4.1 (EL) and over 4.2 times higher average precision than MHCflurry (EL), whereas using BA predictors is not sufficient to accurately predict HLA ligands. The metrics obtained in our benchmark have a substantial gap to the values of a perfect model, suggesting that our benchmark explores a more difficult scenario than the validation schemes typically proposed by other groups, which we think might better reflect the biological situation these computational models are applied to.

## Conflicts of interest

MGK is a consultant to Ardigen.

## Acknowledgments

Martin Klatt is a participant in the BIH Charité Clinician Scientist Program funded by the Charité – Universitätsmedizin Berlin, and the Berlin Institute of Health at Charité (BIH). ARDisplay-I was developed with funding through the project ”Creating an innovative AI-based (Artificial Intelligence) IN SILICO TECHNOLOGY TCRact to launch a NEW SERVICE for designing and optimizing T-cell receptors (TCR) for use in cancer immunotherapies” co-funded by the European Regional Development Fund (ERDF) as part of Smart Growth Operational Programme 2014-2020.

## Author contributions

MK and AB contributed equally to the study. SM, PS, RS, JK, and MK wrote the manuscript. SM and BZ performed the experiments. PS, PB, BKJ, MJ, ASD, RS, JK, and AB developed the ARDisplay-I model. VMP set up and performed the benchmarking analysis of ARDisplay-I and other methods. All authors read and approved the manuscript.

---

**Suppl. Fig. 1. Effect of various conditions of acetonitrile on isolation efficiency of modified MHC ligands. (A-C)** The relative amount of modified MHC ligands compared to the total number of MHC ligands per sample. (A) phosphorylated MHC ligands (B) cysteinylated MHC ligands (C) oxidized MHC ligands). **(D-F)** Venn diagrams showing the overlap of unique modified MHC ligands in different ACN conditions.

**Suppl. Fig. 2. Changes in unique MHC ligands by MHC allele. (A-C)** Absolute numbers of MHC ligands for JJN3, LP-1, and Nalm-6 cells assigned to their expressed MHC alleles as determined by netMHCpan 4.1. %rank of 2 was used as a cut-off for binders.

## Materials and Methods

### Cell lines

LP-1, JJN3 and Nalm-6 cells were maintained at Charité-University Medicine Berlin. MHC typings were retrieved either from published data (JCI insight) or the TRON cell line repository (Citation).

All cell lines were maintained in RPMI media and supplemented with 10% FBS and 2 mM glutamine.

### Immunopurification of MHC class I ligands

For immunopurification suspension cells were harvested through direct resuspension, adherent cell lines after incubating for 15 min with CellStripper solution (Corning™, Cat# 25056CI). Harvested cells were pelleted and washed three times in ice-cold sterile PBS (xxx). For experiments with BV173, AML14, and JMN cell lines 50*10^6^ cells were used, for experiments with LP-1, JJN3, and Nalm-6 cell lines 150*10^6^ cells were used. Cells were lysed in 7.5 ml of 1% CHAPS (Sigma-Aldrich, Cat# C3023) dissolved in PBS and supplemented with protease inhibitors (cOmplete, Cat# 11836145001). Cell lysis was performed for 1 hour at 4°C, lysates were spun down for 1 hour with 20,000 g at 4°C, and supernatant fluids were isolated. Affinity columns were prepared as follows: 40 mg of Cyanogen bromide-activated-Sepharose 4B (Sigma-Aldrich, Cat# C9142) were activated with 1mM hydrochloric acid (Sigma-Aldrich, Cat# 320331) for 30 min. Subsequently, 1 mg of W6/32 antibody (BioXCell, Cat #BE0079) was coupled to sepharose in the presence of binding buffer (150mM sodium chloride, 50 mM sodium bicarbonate, pH 8.3; sodium chloride: Sigma-Aldrich, Cat# S9888, sodium bicarbonate: Sigma-Aldrich, Cat#S6014) for at least 2 hours at room temperature. Sepharose was blocked for 1 h with glycine (Sigma-Aldrich, Cat# 410225) and washed 3 times with PBS.

Supernatants of cell lysates were run over the columns through peristaltic pumps with a 1 ml/min flow rate overnight in a cold room. Affinity columns were washed with PBS for 30 min, water for 30 min, then run dry, and MHC complexes were subsequently eluted five times with 200 μl 1% trifluoracetic acid (TFA, Sigma/Aldrich, Cat# 02031). The TFA eluates were pooled and then split in as many portions as settings were investigated (either 3 portions with 30%, 40%, and 50% ACN settings and 5 portions with 5%, 20%, 35%, 50%, and “mix” settings).

For separation of MHC ligands and their MHC complexes C18 columns (Sep-Pak C18 1 cc Vac Cartridge, 50 mg Sorbent per Cartridge, 37-55 μm Particle Size, Waters, Cat# WAT054955) were prewashed with 80% acetonitrile (ACN, Sigma-Aldrich, Cat# 34998) in 0.1% TFA and equilibrated with two washes of 0.1% TFA. Samples were loaded, washed again with 0.1% TFA and eluted in 400 μl of either 30%, 40%, or 50% ACN in 0.1%TFA for BV173, AML14, and JMN cells or for LP-1, JJN3, and Nalm-6 cells with 4x200 ul of either 5%, 20%, 35% and 50% ACN in 0.1%TFA or 200ul each of 5%, 20%, 35% and 50% ACN in 0.1%TFA for the “mix” setting. The sample volume was reduced by vacuum centrifugation for mass spectrometry analysis.

### Solid Phase Extractions (SPE)

In-house C18 mini columns were prepared as follows: for SPE of one sample two small disks of C18 material (1mm in diameter) were punched out from CDS Empore™ C18 disks (Fisher Scientific, Cat# 13-110-018) and transferred to the bottom of a 200 µl Axygen pipette tip (Fisher Scientific, Cat# 12639535). Columns were washed once with 100 µl 80%ACN/0.1%TFA and equilibrated with 3 times 100 µl 1%TFA. All fluids were run through the column by centrifugation in mini tabletop centrifuges and eluates were collected in Eppendorf tubes. Then, dried samples were resuspended in 100 µl 1%TFA and loaded onto the columns, washed twice with 100 µl 1%TFA, ran dry, and eluted with 50 µl 80%ACN/0.1% TFA. Again, the sample volume was reduced by vacuum centrifugation.

### LC-MS/MS analysis of MHC ligands

Samples were analyzed by high resolution/high accuracy LC-MS/MS (Lumos Fusion, Thermo Fisher). Peptides were separated using direct loading onto a packed-in-emitter C18 column (75um ID/12cm, 3 μm particles, Nikkyo Technos Co., Ltd. Japan). The gradient was delivered at 300 nl/min increasing linear from 2% Buffer B (0.1% formic acid in 80% acetonitrile) / 98% Buffer A (0.1% formic acid) to 30% Buffer B / 70% Buffer A, over 70 minutes. MS and MS/MS were operated at resolutions of 60,000 and 30,000, respectively. Only charge states 1, 2, and 3 were allowed. 1.6 Th was chosen as the isolation window and collision energy was set at 30%. For MS/MS, the maximum injection time was 100ms with an AGC of 50,000.

### Mass spectrometry data processing

Mass spectrometry data were processed using Byonic software (version 4.5.2, Protein Metrics, Palo Alto, CA) through a custom-built computer server. Mass accuracy for MS1 was set to 10 ppm and 20 ppm for MS2, respectively. Digestion specificity was defined as unspecific and only precursors with charges 1,2, 3, and up to 2 kDa were allowed. Protein FDR was disabled to allow a complete assessment of potential peptide identifications. Oxidation of methionine, phosphorylation of serine, threonine, and tyrosine as well as cysteinylation of cysteine were set as variable modifications for all samples. Samples were searched against the UniProt Human Reviewed Database with common contaminants added. Peptides were selected with a minimal log prob value of 1.3 indicating p-values for PSMs of <0.05 and duplicates removed.

### Assignment of peptide sequences to MHC alleles

To assign peptides that passed the MS quality filters described above to their MHC alleles which they most likely bind to, we used the netMHCpan 4.1 algorithm with default settings12, and the ARDisplay-I presentation model. No binding affinity predictions were enabled. Therefore, all peptides with %ranks below 2 were considered binders for netMHCpan and above the presentation probability of 0.5 for ARDisplay-I.

### Software and Statistics

All graphs except Venn diagrams were drawn with GraphPad Prism 7. For statistics built-in analyses from GraphPad Prism were used. One-way ANOVA tests with Friedmann’s multiple comparison tests were used for comparing GRAVY scores. Venn diagrams were prepared using the Venn diagrams tool by the University of Gent. (https://bioinformatics.psb.ugent.be/webtools/Venn/)

### Training of ARDisplay-I, ie. HLA ligand presentation model

A curated dataset containing peptides presented by class I HLAs on the surface of host cells was extracted from more than 20 publicly available datasets from selected publications. The presence of each HLA ligand was experimentally confirmed by MS experiments provided by each group. All peptides were of human origin and were presented on the surface of either mono-allelic human cell lines or multi-allelic data obtained from healthy donors or cancer patients. Synthetic negative data (ie. non-presented peptides) were prepared based on human proteome (GRChg38, release 98). Only peptides containing 8-11 naturally occurring amino acids were taken into consideration.

The current version of ARDisplay-I is a classifier combining a custom approach to multi-allelic data incorporation (ie. by extracting all possible pHLA pairs, generating a model prediction for each, applying filtering based on BA predictions from MHCflurry and re-aggregating the results) together with BERT^28^/TAPE^29^ embeddings and simple MLP (Multi-Layer Perceptron) as last layers, and model fine-tuning to HLA eluted ligands prediction. The aforementioned BERT (bidirectional encoder representations from transformers) is an NLP family of models based on transformer architecture. Such a model was applied to selected tasks involving protein biology - see TAPE (Tasks Assessing Protein Embeddings). One of the TAPE models pre-trained with a semi-supervised approach used for protein representation learning, was used as an initial step for the training of ARDisplay-I. Our model outputs the presentation probability of a given pHLA pair. Moreover, ARDisplay-I can be used to predict whether a peptide will be displayed via any of the patients’ HLA molecules. To ensure that ARDisplay-I can make predictions on a vast set of known HLA alleles we used the pseudo-sequences provided in the PUFFIN model^30^ to encode the HLA types. Such pseudo-sequences describe those amino acids from the HLA that form a binding groove and have a chance to interact with the peptide that they bind.

ARDisplay-I was implemented in Python 3.9 using the PyTorch v.1.13.1 library that enables performing fast operations on tensors and neural networks with GPU acceleration. Additionally, we have used PyTorch Lightning v.2.0.0 for the maintenance and training of our model. GPU-based computations were done on a machine equipped with NVIDIA Ampere A100 GPU card with CUDA® 8.6 architecture, 640 Tensor Cores, 6,912 CUDA® Cores, and 40 GB HBM2 GPU Memory and using cudnn 8.5.0 and cudatoolkit 11.7.99. Standard Python libraries for data analysis and machine learning were used, inter alia, scikit-learn, pandas, numpy, and matplotlib with seaborn.

### Metrics used for benchmarking

Precision-recall curve (PR curve) is a standard curve for binary classification model evaluation. It is especially useful for very imbalanced data and visualizes a trade-off between precision (the proportion of positive predicted samples that are true positives) and recall (the fraction of all positives detected by the model). Average precision (AP) is a weighted mean of precision at each threshold, ie. a proxy to the area under the precision-recall curve (AUPRC).

Precision (also positive predictive value, PPV) is a metric that describes the percentage of positively classified cases in binary classification that are true positives. In the case of presentation models, PPV provides information on what fraction of the pHLA pairs, labeled by the model as presented, actually turned out to be presented in the experimental setup. Typically, PPV estimation requires defining a threshold above which an observation is classified as positive. For example, the score for PPV top-100 describes how many true positive observations can be expected by testing in the lab the 100 pHLA pairs with the highest ranks as determined by the model predictions.

## Footnotes

1 tumor-infiltrating lymphocytes

2 human leukocyte antigen, also known as human version of MHC

## Notes

https://huggingface.co/spaces/ardigen/ardisplay-i

