## Supplementary figures and images for "Identification of tumor-specific MHC ligands through improved biochemical isolation and incorporation of machine learning"

### Supplementary Figure 1

Suppl. Figure 1

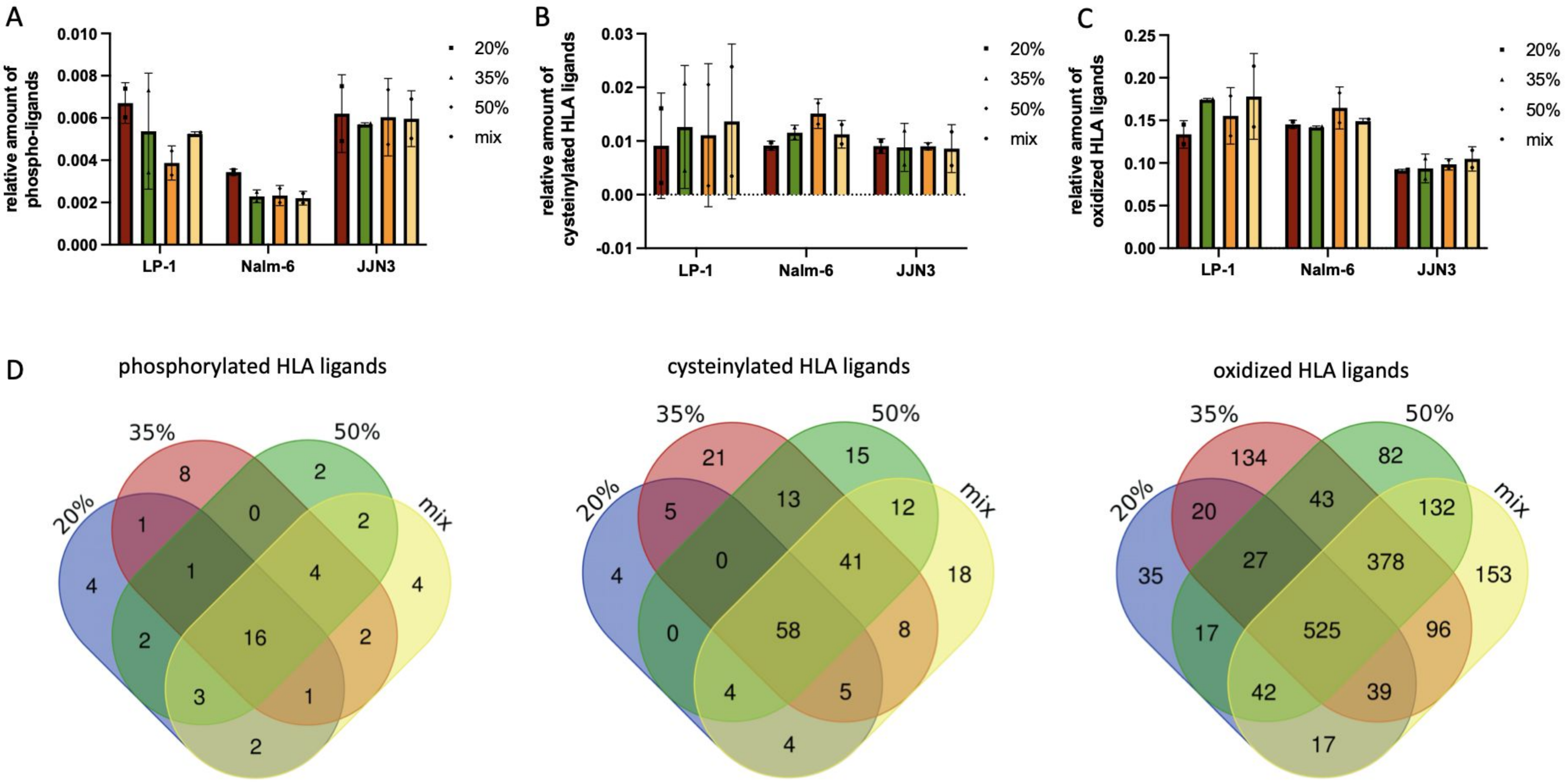

### Supplementary Figure 2

Suppl. Figure 2

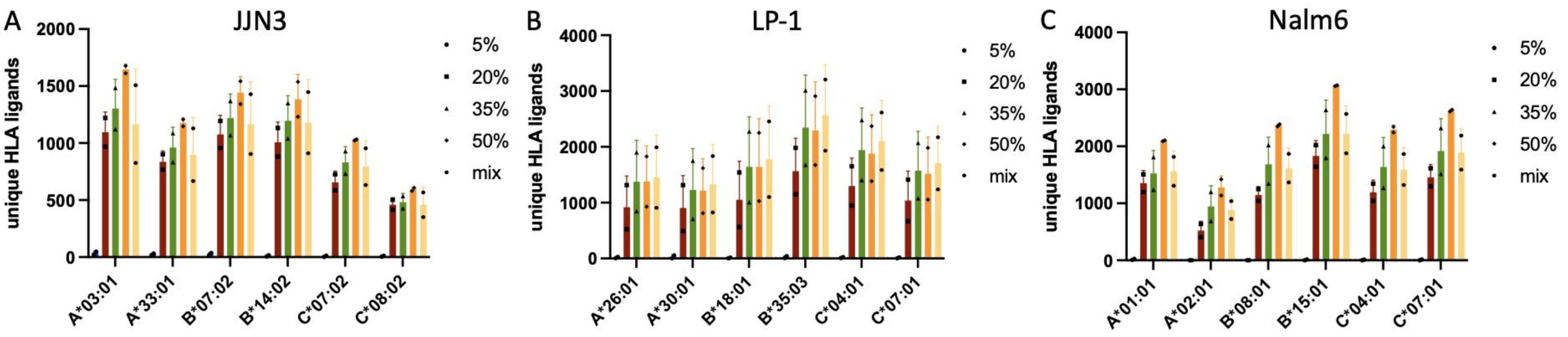
